# Genome-wide association study of grain iron and zinc concentrations in a diverse CIMMYT wheat panel across contrasting moisture environments

**DOI:** 10.64898/2026.08.06.743196

**Authors:** Kai Yuan, Yi Dai, Zerihun Tadesse Tarekegn, Lahu Lu, Xiaofei Ma, Velu Govindan

## Abstract

Micronutrient deficiencies remain a major public-health challenge, and genetic improvement of grain iron and zinc concentrations in wheat offers a sustainable biofortification strategy. We evaluated 563 CIMMYT advanced wheat lines and six checks under restricted irrigation (two irrigations) and well-watered conditions (five irrigations) at Ciudad Obregón, Mexico. Grain iron and zinc concentrations were quantified by energy-dispersive X-ray fluorescence spectrometry. Best linear unbiased estimates were calculated, and genome-wide association analyses were conducted using 8,687 high-quality single-nucleotide polymorphisms and a multi-locus mixed model that accounted for population structure. Five marker-trait associations were detected in at least two datasets. For grain zinc concentration, *S3B_811421507* was detected under both irrigation regimes and in the combined analysis, whereas *S3B_814372642* was detected under well-watered conditions and in the combined analysis. For grain iron concentration, *S2B_73321328* and *S4B_20679079* were associated with variation under restricted irrigation and in the combined analysis, while *S2B_72162723* was detected under well-watered conditions and in the combined analysis. Individual loci explained 0.34% to 3.36% of phenotypic variance, consistent with the quantitative inheritance of grain micronutrient concentration. The identified alleles provide candidate targets for validation and marker-assisted biofortification breeding, while their environment-dependent effects emphasize the need to evaluate micronutrient traits across contrasting moisture conditions.

## Introduction

Micronutrient deficiency, often described as hidden hunger, is a major global public-health challenge. More than two billion people have inadequate dietary intake of iron (Fe) or zinc (Zn), particularly in regions where cereal-based diets predominate [1, 2, 3]. Iron and zinc are essential for normal physiological function, and chronic deficiencies contribute to anemia and impaired immune function [4]. Wheat (*Triticum aestivum L*.) supplies approximately one-fifth of global dietary calories and protein [5] and is also an important source of micronutrients [6, 7]. However, many modern high-yielding cultivars have relatively low concentrations of Fe and Zn in the edible grain [8, 9, 10]. Genetic biofortification is therefore a cost-effective and sustainable approach for increasing micronutrient intake through commonly consumed foods [11], and marker-assisted selection can accelerate the introgression of favorable alleles into breeding populations [12].

Grain micronutrient accumulation is a complex quantitative process controlled by many loci and influenced by environmental conditions, including soil properties, water availability, and temperature [13]. Quantitative trait locus mapping and genome-wide association studies have identified loci for grain zinc concentration (GZnC) and grain iron concentration (GFeC) across all wheat chromosomes [7, 14, 15, 16]. Some loci affect both traits, suggesting pleiotropy or tight linkage [17]. A previous study of 330 wheat accessions identified 39 loci associated with GZnC and GFeC [18]. However, the stability of these associations across moisture regimes remains insufficiently characterized.

We evaluated 563 CIMMYT wheat advanced breeding lines under restricted and normal irrigation and used genotyping-by-sequencing markers to investigate the genetic basis of GFeC and GZnC. The objectives were to identify marker-trait associations that were reproducible across moisture environments, estimate their allelic effects, and identify candidate loci for validation and use in wheat biofortification breeding.

## Materials and methods

### Plant materials and experimental design

The association panel comprised 563 CIMMYT advanced wheat breeding lines and six checks (S1 Table).

The field experiment was conducted at the Norman E. Borlaug Experimental Station (CENEB), Ciudad Obregón, Mexico, during the 2023-2024 cropping season. The panel was evaluated under two irrigation regimes: two irrigations (2IR), representing restricted irrigation, and five irrigations (5IR), representing well-watered conditions. An alpha-lattice design with two replications was used. Each entry was sown in a six-row plot 2.8 m long with 0.30 m between rows, giving a plot area of 4.48 m2. Phenotypic datasets were designated 2IRFe, 2IRZn, 5IRFe, and 5IRZn. Crop management followed standard recommendations [10]. Before sowing, 25 kg ha^-1^ ZnSO4·7H2O, 50 kg ha^-1^ nitrogen, and 80 kg ha^-1^ P_2_O_5_ were applied. An additional 100 kg ha^-1^ nitrogen was top-dressed approximately 30 days after sowing. Standard chemical controls were used to manage pests and diseases.

### Quantification of grain iron and zinc concentrations

Twenty-five to thirty spikes were harvested manually from each plot using procedures designed to minimize metallic and environmental contamination. A 20-g grain sample from each plot was cleaned to remove cracked kernels, chaff, and other impurities. Grain Fe and Zn concentrations (mg kg-1) were measured using an X-Supreme 8000 energy-dispersive X-ray fluorescence spectrometer (Oxford Instruments, Abingdon, UK), a validated non-destructive method for wheat grain analysis [19]. Each sample was measured three times, and the mean was used in subsequent analyses.

### Statistical analysis of phenotypic data

Phenotypic data were analyzed using META-R version 6.04 [20]. Variance components and best linear unbiased estimates (BLUEs) were obtained by restricted maximum likelihood using the lmer function in the R package lme4.

Broad-sense heritability (H^2^) within each irrigation treatment was estimated from the genetic and residual variance components, accounting for two replications. Pearson correlation coefficients were calculated among traits and environments. Statistical analyses were conducted in R and IBM SPSS Statistics version 25.0.

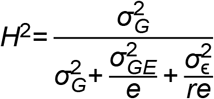

### Genomic DNA extraction and SNP genotyping

Genomic DNA was extracted from two to three young leaves collected from 14-day-old seedlings using BioSprint 96 Plant Kits (Qiagen, Hombrechtikon, Switzerland). Genotyping-by-sequencing libraries were prepared in 96-plex format after digestion with PstI and MspI [21] and sequenced on Illumina platforms at the Wheat Genetics Resource Center, Kansas State University, USA. Single-nucleotide polymorphisms (SNPs) were called with the TASSEL 5.2.40 GBS v2 pipeline [22] and aligned to the Chinese Spring reference genome (IWGSC RefSeq v1.0; International Wheat Genome Sequencing Consortium 2018). The minimum tag count was set to five. SNPs with minor allele frequency below 0.05 or more than 50% missing data were removed. All lines had less than 20% missing data. After filtering, 8,687 SNPs were retained (S2 Table).

### Population structure and linkage disequilibrium analysis

Population structure was evaluated with ADMIXTURE version 1.3.0 for K = 1-10. Principal component analysis and the genomic relationship matrix were calculated in GAPIT 3.0; kinship was estimated using the VanRaden method. Linkage disequilibrium (LD), measured as pairwise r2, was calculated with PopLDdecay [23]. LD decay distance was defined as the physical distance at which r2 declined to half of its maximum value.

### Genome-wide association analysis

Genome-wide association analyses were performed for 2IR, 5IR, and BLUE datasets using the multi-locus mixed model implemented in GAPIT [7]. The first three principal components were included as fixed covariates to account for population structure. Associations with - log10(P) > 3 were retained. An association was considered stable when detected in at least two datasets. The phenotypic variance explained by each lead SNP was estimated following Shim et al. [24].

Manhattan and quantile-quantile plots were generated with the R package CMplot.

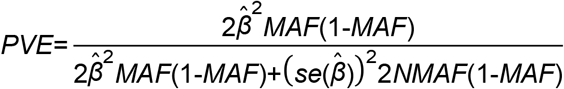

## 3. Results

### Phenotypic and genetic variation across moisture environments

Both GFeC and GZnC showed continuous variation under the two irrigation regimes (Table 1; Fig 1). Mean GFeC was 36.34 mg kg^-1^ under 2IR and 35.79 mg kg^-1^ under 5IR, with broad-sense heritabilities of 50.61% and 55.83%, respectively. Mean GZnC was 45.75 mg kg^-1^ under 2IR and 45.89 mg kg^-1^ under 5IR, with heritabilities of 60.86% and 69.99%, respectively. Thus, GZnC showed greater phenotypic variation and higher heritability than GFeC in both environments.

**Table 1.** Distribution of 563 wheat Fe and Zn values across two treatments.

| Traits | Env. | Min | Max | Mean | SD | CV (%) | H<sup>2</sup> (%) |
| --- | --- | --- | --- | --- | --- | --- | --- |
| GFeC | 2IR | 26.40 | 47.40 | 36.34 | 3.15 | 8.66 | 50.61 |
|  | 5IR | 27.00 | 51.10 | 35.79 | 3.00 | 8.38 | 55.83 |
|  | BLUE | 28.60 | 47.70 | 35.98 | 2.06 | 5.72 | - |
| GZnC | 2IR | 28.40 | 64.90 | 45.75 | 4.91 | 10.74 | 60.86 |
|  | 5IR | 30.20 | 65.40 | 45.89 | 5.12 | 11.15 | 69.99 |
|  | BLUE | 37.30 | 56.60 | 46.19 | 3.28 | 7.11 | - |

**Fig 1.**
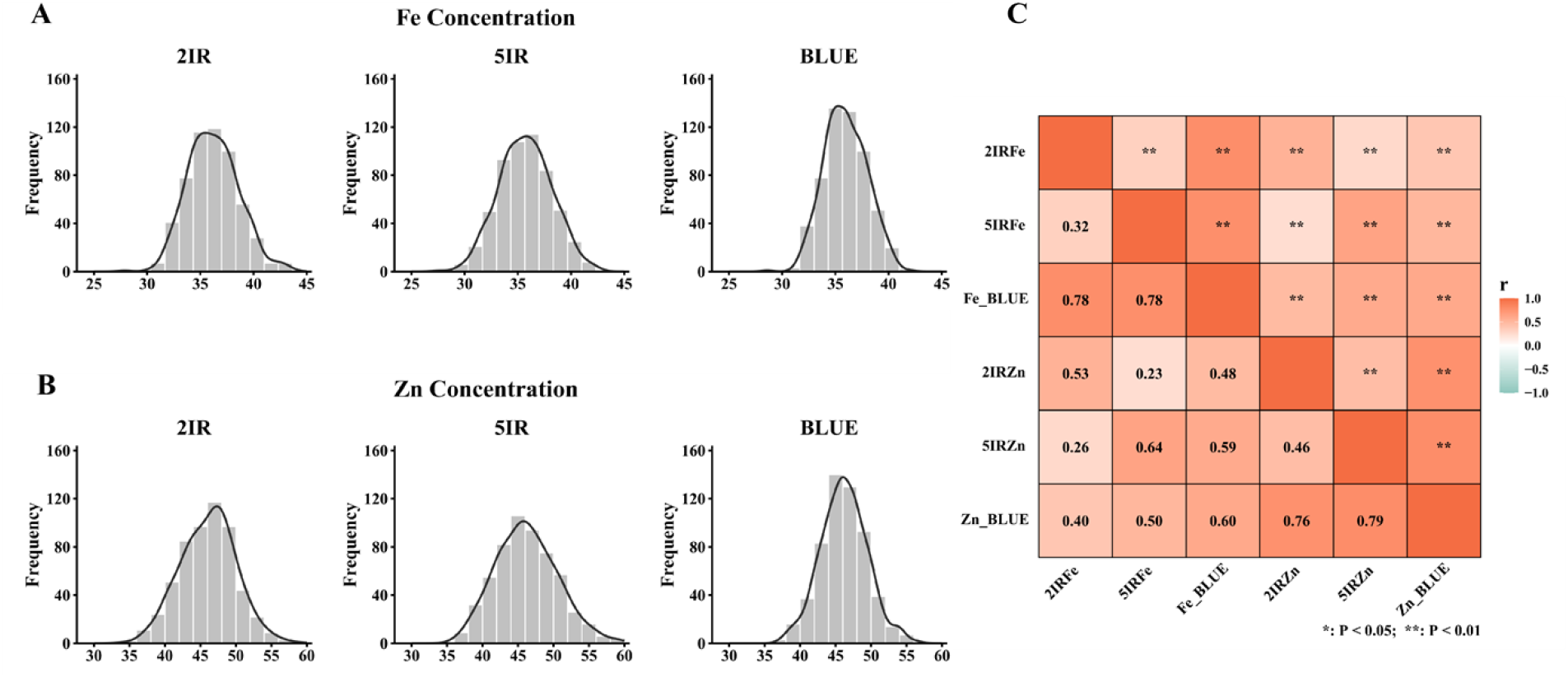
Phenotypic distribution and correlation analysis of grain iron (Fe) and zinc (Zn) concentrations in the wheat population. **A**. Frequency distribution of grain Fe concentration under 2IR, 5IR, and BLUE environments. **B**. Frequency distribution of grain Zn concentration under 2IR, 5IR, and BLUE environments. The black curves represent the normal distribution fit. **C**. Pearson correlation heatmap of grain Fe and Zn concentrations across different environments. Values in the lower triangle are Pearson correlation coefficients (*r*). The upper triangle displays the corresponding significance levels (*P* < 0.05, *P* < 0.01).

Distributions of GFeC and GZnC were approximately continuous in each environment and for BLUEs (Fig 1A and 1B). Measurements of the same trait across irrigation regimes were positively correlated (r = 0.32 for Fe and r = 0.46 for Zn; P < 0.01). Fe_BLUE and Zn_BLUE were also positively correlated (Fig 1C).

### Population structure and linkage disequilibrium

Of the 8,687 retained SNPs, 3,614 (41.60%) mapped to the A genome, 4,213 (48.50%) to the B genome, and 860 (9.90%) to the D genome. Chromosome 2B contained the most markers (903; 10.39%), whereas chromosome 4D contained the fewest (44; 0.51%) (S5 Table). ADMIXTURE supported three genetic subgroups (Fig 2A). LD decayed to half of its maximum at approximately 3.1 Mb, 3.9 Mb, and 3.5 Mb in the A, B, and D genomes, respectively (Fig 2B).

**Fig 2.**
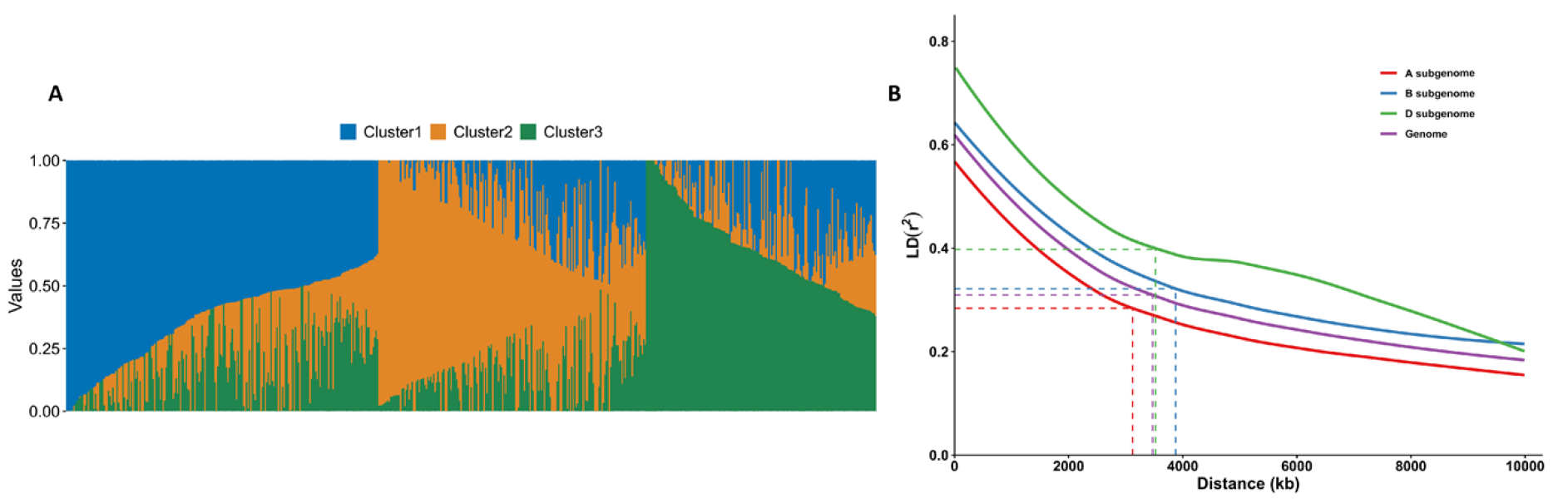
**Population structure and linkage disequilibrium analysis based on the SNPs. A The ancestral fractions estimated for each varieties using software ADMIXTURE for K = 3, different colors represent different clusters. B The LD decay for the A, B, and D subgenomes and the entire genome of bread wheat**

### Genome-wide association analysis

Three stable associations were detected for GFeC (Table 2; Fig 3). The marker *S2B_72162723* was detected in the 5IR and BLUE datasets. *S2B_73321328* and *S4B_20679079* were detected in the 2IR and BLUE datasets and explained up to 2.66% of phenotypic variance.

**Table 2.** Significant single nucleotide polymorphisms (SNPs) associated with GFeC and GZnC.

| Trait | SNP | Chr. | Pos. | Env. | Alleles | $-\log_{10}(p)$ | PVE |
| --- | --- | --- | --- | --- | --- | --- | --- |
| Fe | S2B_72162723 | 2B | 72162723 | 5IR, BLUE | C/T: C | 3.08-3.11 | 0.34-0.59 |
|  | S2B_73321328 | 2B | 73321328 | 2IR, BLUE | C/G: C | 3.16 | 2.01 |
|  | S4B_20679079 | 4B | 20679079 | 2IR, BLUE | C/T: T | 3.70-4.06 | 2.39-2.66 |
| Zn | S3B_811421507 | 3B | 811421507 | 2IR, 5IR, BLUE | G/A: A | 3.09-5.01 | 1.95-3.36 |
|  | S3B_814372642 | 3B | 814372642 | 5IR, BLUE | A/G: G | 3.23-3.64 | 2.05-2.35 |

**2IR:** 2 irrigation; **5IR:** 5 irrigation; **BLUE:** best linear unbiased estimation; **PVE:** phenotypic variance explained (%)

**Fig 3.**
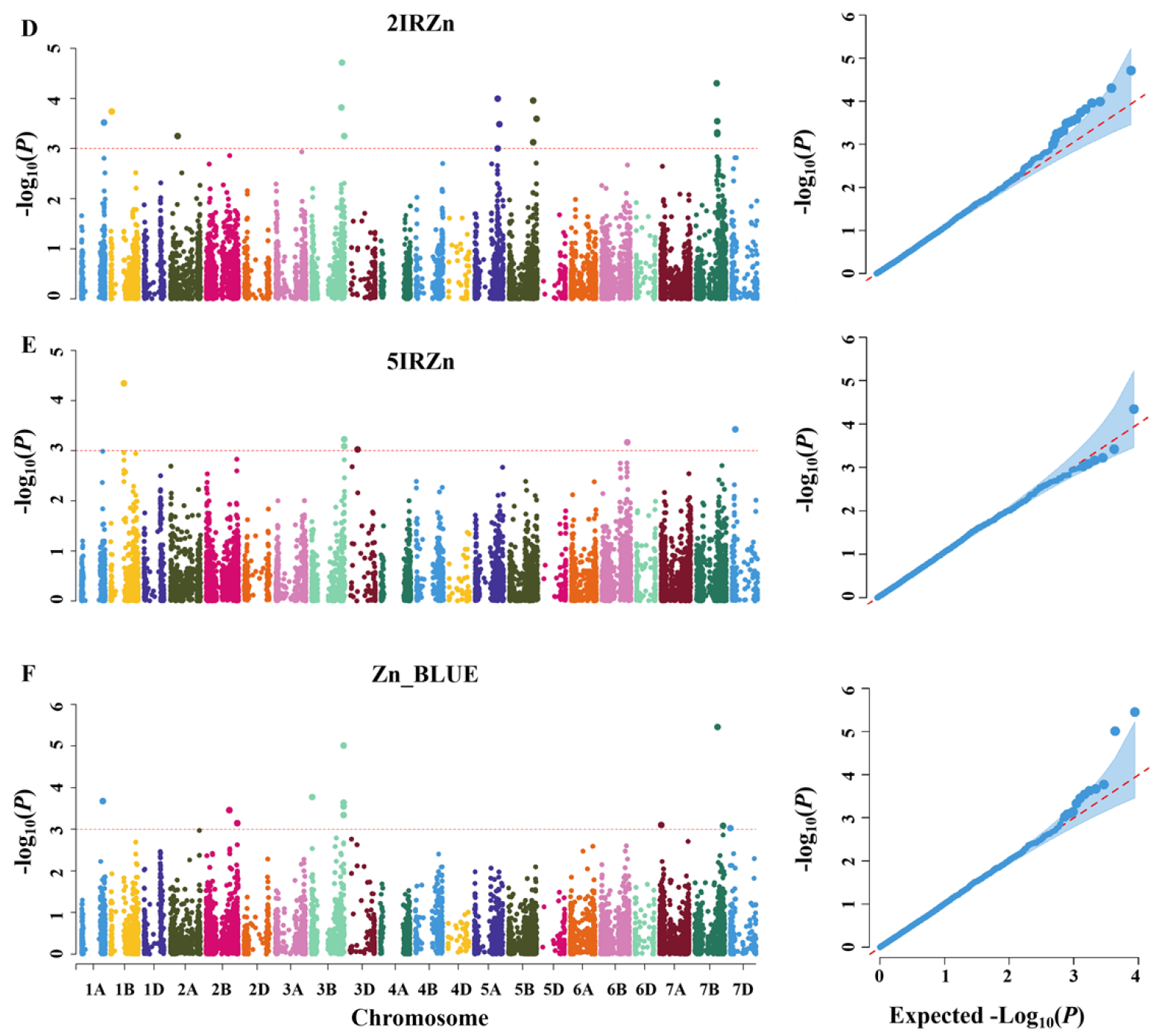
**Manhattan and Q-Q plots for grain Fe and Zn contents based on BLUE values. A: 2IRFe, B: 5IRFe, C: Fe_BLUE, D: 2IRZn, E: 5IRZn, and F: Zn_BLUE. The horizontal dashed lines represent the significance threshold (P <1×10-3).**

Two stable associations were detected for GZnC, *S3B_811421507* was detected in 2IR, 5IR, and BLUE datasets, with a maximum -log10(P) of 5.01 and phenotypic variance explained of 3.36% and *S3B_814372642* was detected in the 5IR and BLUE datasets (Table 2; Fig 3).

### Allelic effects at significant loci

Both stable GZnC loci were located on chromosome 3B (Fig 4D). At S3B_811421507, lines carrying AA had higher GZnC than lines carrying GG under 5IR and for BLUEs (Fig 4E). At S3B_814372642, GG lines had higher GZnC than AA lines under 5IR and for BLUEs, whereas the difference was not significant under 2IR (Fig 4F).

**Fig 4.**
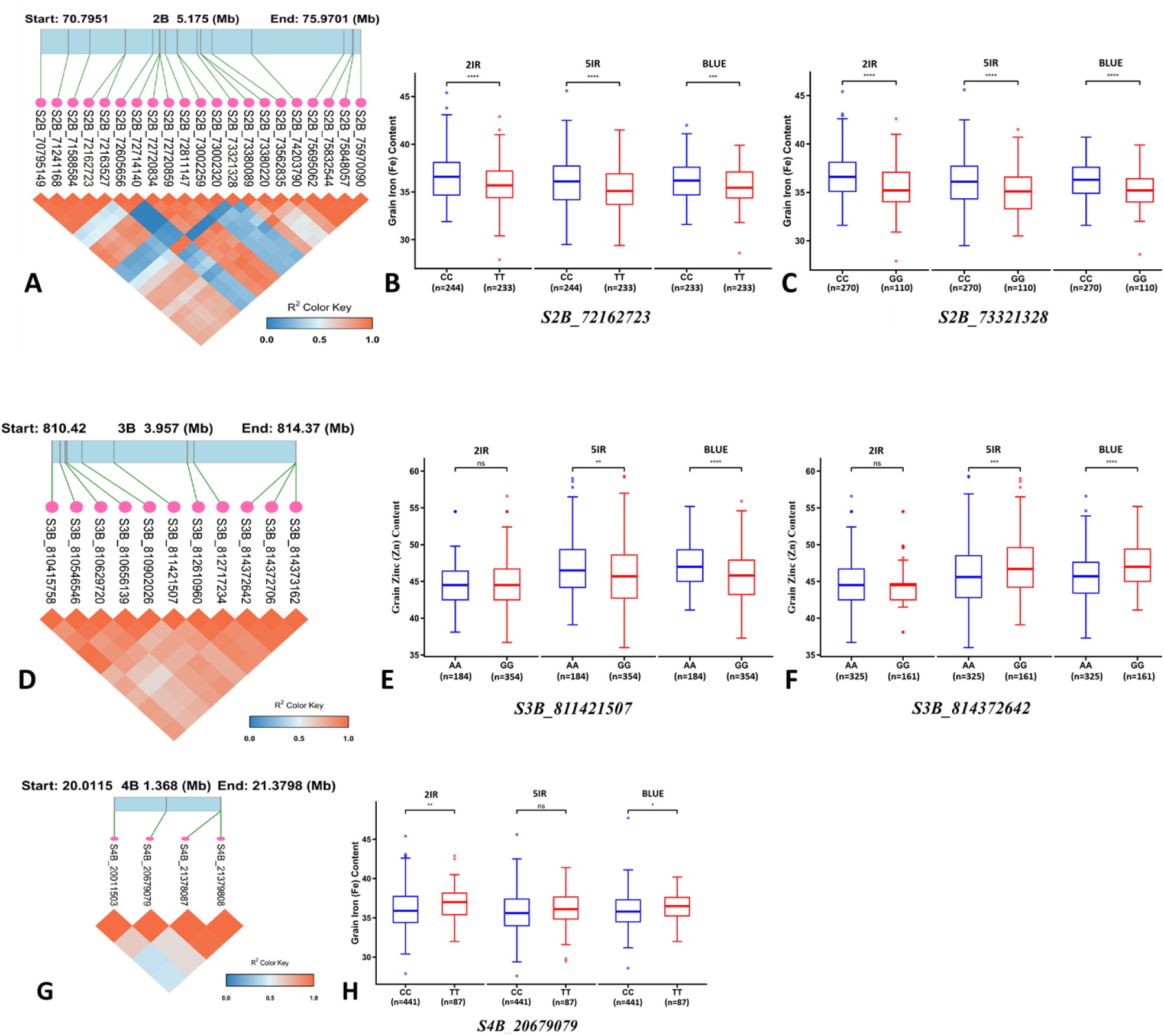
LD block analysis and phenotypic effects of superior alleles for the significant MTAs. **A, D, G:** Local LD heatmaps of the genomic regions on chromosomes 2B, 3B, and 4B, respectively. The color key at the bottom indicates the level of linkage disequilibrium (R^2^ values) between markers, with darker colors representing stronger LD. **B, C, H:** Phenotypic differences in grain iron (Fe) concentration between diverse alleles of the core SNPs on chromosomes 2B and 4B across 2IR, 5IR, and BLUE datasets. E, F: Phenotypic differences in grain zinc (Zn) concentration between distinct alleles of the core SNPs on chromosome 3B. (The “n” below each boxplot indicates the number of accessions carrying the respective allele). Statistical significance was determined using a two-tailed Student’s t-test (ns, not significant; * *P* < 0.05; \*\**P* < 0.01; *** *P* < 0.001; **** *P* < 0.0001).

## Discussion

Positive correlations between grain Fe and Zn concentrations can facilitate simultaneous improvement of both traits. Previous studies have attributed such correlations to linked genes controlling micronutrient uptake and translocation or to pleiotropic regulators [7, 25, 26]. The Gpc-B1 locus, for example, affects grain protein, Zn, and Fe concentration by promoting nutrient remobilization during senescence [27, 28]. In this study, Fe and Zn were positively correlated across irrigation regimes and BLUE datasets. These results support joint selection for both micronutrients, although the moderate correlations also indicate that each trait retains substantial independent variation.

The five stable associations explained 0.34%-3.36% of phenotypic variance, consistent with the minor-effect architecture expected for grain micronutrient concentration [10, 29]. No locus was significantly associated with both Fe and Zn. This may reflect true trait-specific effects or the limited marker density of the GBS dataset, which reduces power to detect small-effect or pleiotropic loci. The chromosome 3B region identified here is close to S3B_813450132, previously associated with GZnC in synthetic hexaploid wheat [30]. However, local LD analysis suggests that *S3B_811421507* and *S3B_813450132* are not within the same strong LD block and may represent independent signals.

A previously reported GFeC QTL, QGFezx.caas-4BS, spans 22.3-36.0 Mb on chromosome 4B [31]. *S4B_20679079*, detected at 20.7 Mb in the present study, is near this interval. Given the estimated 3.9-Mb LD decay distance in the B genome, the two signals may represent the same or closely linked loci; additional fine mapping and functional validation are required.

The two chromosome 2B associations, *S2B_72162723 and S2B_73321328*, were not located in the same strong local LD block. This pattern may result from low marker density, elevated local recombination, or two independent loci [32, 33]. The environment-dependent effects observed for the Fe and Zn loci highlight genotype-by-environment interactions and indicate that marker validation should include both restricted and well-watered conditions before deployment in breeding.

## Conclusions

Genome-wide association analysis of 563 CIMMYT wheat lines identified five associations for grain Fe and Zn concentrations that were detected in at least two moisture-related datasets. The marker *S3B_811421507* was the most consistently detected GZnC association, whereas the GFeC associations on chromosomes 2B and 4B showed irrigation-dependent effects. Because all loci explained relatively small proportions of phenotypic variance, they should be validated in independent populations and converted to breeder-friendly KASP markers before routine use. The results provide candidate genetic resources for wheat biofortification and demonstrate the importance of evaluating micronutrient traits across contrasting moisture environments.

The authors have declared that no competing interests exist.

## Author contributions

Conceptualization: KY, VG. Data curation: YD, ZTT, LL, XM. Formal analysis: KY, YD, LL, XM. Investigation: KY, YD, ZTT, LL, XM. Methodology: KY, YD, ZTT, LL, XM, VG. Supervision: VG. Writing - original draft: KY. Writing - review & editing: all authors.

## Acknowledgments

All data underlying the findings should be deposited in a public repository or supplied as Supporting Information before acceptance. Replace this sentence with the repository name and persistent identifier, or state that all relevant data are within the manuscript and its Supporting Information files.

## Data availability

All data underlying the findings are supplied as Supporting Information.

## Supporting information captions

S1 Table. List of 563 CIMMYT advanced wheat breeding lines and six checks evaluated in the association panel.

S2 Table. Genotyping-by-sequencing marker dataset retained after quality filtering.

S3 Table. Genome-wide association results for grain iron and zinc concentrations across irrigation regimes and BLUE datasets.

S4 Table. Phenotypic values for grain iron and zinc concentrations under 2IR and 5IR conditions.

S5 Table. Distribution of retained SNP markers across wheat chromosomes and subgenomes.

## References

1. Harding KL, Aguayo VM, Webb P. 2018; Hidden hunger in South Asia: a review of recent trends and persistent challenges. Public Health Nutr 21(4):785–795. doi: 10.1017/S1368980017003202

2. Liu ZH, Wang HY, Wang XE, Zhang GP, Chen PD, Liu DJ. 2006; Genotypic and spike positional difference in grain phytase activity, phytate, inorganic phosphorus, iron, and zinc contents in wheat (triticum aestivum L.). J Cereal Sci 44(2):212–219. doi: 10.1016/j.jcs.2006.06.001

3. Stein AJ, Qaim M. 2007; The human and economic cost of hidden hunger. Food Nutr Bull 28(2):125–34. 10.1177/156482650702800201

4. Black RE, Victora CG, Walker SP, Bhutta ZA, Christian P, de Onis M et al. 2013; Maternal and child undernutrition and overweight in low-income and middle-income countries. Lancet 382(9890):427–451. 10.1016/S0140-6736(13)60937-X

5. Shiferaw B, Smale M, Braun HJ, Duveiller E, Reynold M, Muricho G. 2013; Crops that feed the world 10. Past successes and future challenges to the role played by wheat in global food security. Food Sec 5:291–37.10.1007/s12571-013-0263-y

6. Ludwig Y, Slamet-Loedin IH. 2019; Genetic Biofortification to Enrich Rice and Wheat Grain Iron: From Genes to Product. Front Plant Sci 10:833. doi: 10.3389/fpls.2019.00833

7. Wang J, Zhang Z. 2021; GAPIT Version 3: Boosting Power and Accuracy for Genomic Association and Prediction. Genomics Proteomics Bioinformatics 19(4):629–640. doi: 10.1016/j.gpb.2021.08.005

8. Alchoubassi G, Aszyk J, Pisarek P, Bierla K, Ouerdane L, Szpunar J, Lobinski R. 2018; Advances in mass spectrometry for iron speciation in plants. TrAC Trends in Analytical Chemistry 104:77–86. doi: 10.1016/j.trac.2017.11.006

9. Hui X, Luo L, Chen Y, Palta JA, Wang Z. 2025; Zinc agronomic biofortification in wheat and its drivers: a global meta-analysis. Nat Commun 16(1):3913. doi: 10.1038/s41467-025-58397-y

10. Velu G, Singh RP, Huerta J, Guzmán C. 2017; Genetic impact of Rht dwarfing genes on grain micronutrients concentration in wheat. Field Crops Res 214:373–377. doi: 10.1016/j.fcr.2017.09.030

11. Bouis HE, Hotz C, McClafferty B, Meenakshi JV, Pfeiffer WH. 2011; Biofortification: a new tool to reduce micronutrient malnutrition. Food Nutr Bull 32(1 Suppl):S31–40. doi: 10.1177/15648265110321S105

12. Dhaliwal SS, Ram H, Shukla AK, Mavi GS. 2019; Zinc biofortification of bread wheat, triticale, and durum wheat cultivars by foliar zinc fertilization. Journal of Plant Nutrition 42(8):813–822. doi: 10.1080/01904167.2019.1584218

13. Peleg Z, Cakmak I, Ozturk L, Yazici A, Jun Y, Budak H et al. 2009; Quantitative trait loci conferring grain mineral nutrient concentrations in durum wheat x wild emmer wheat RIL population. Theor Appl Genet 119(2):353–69. doi: 10.1007/s00122-009-1044-z

14. Gupta PK, Balyan HS, Sharma S, Kumar R. 2021; Biofortification and bioavailability of Zn, Fe and Se in wheat: present status and future prospects. Theor Appl Genet 134(1):1–35. doi: 10.1007/s00122-020-03709-7

15. Tsonev S, Dragov R, Taneva K, Christov NK, Bozhanova V, Todorovska EG. 2024; Genome-Wide Association Studies of Agronomic and Quality Traits in Durum Wheat. Agriculture 14(10):1743. doi: 10.3390/agriculture14101743

16. Zhang Z, Lan C, Singh RP, Govindan V. 2026; Genetic dissection of grain zinc, iron, and yield traits in a CIMMYT bread wheat mapping population to support biofortification breeding. Mol Breed 46(6):56. 10.1007/s11032-026-01663-8

17. Tong J, Sun M, Wang Y, Zhang Y, Rasheed A, Li M et al. 2020; Dissection of Molecular Processes and Genetic Architecture Underlying Iron and Zinc Homeostasis for Biofortification: From Model Plants to Common Wheat. Int J Mol Sci 21(23):9280. doi: 10.3390/ijms21239280

18. Velu G, Singh RP, Crespo-Herrera L, Juliana P, Dreisigacker S, Valluru R. 2018; Genetic dissection of grain zinc concentration in spring wheat for mainstreaming biofortification in CIMMYT wheat breeding. Sci Rep 8(1): 13526. doi: 10.1038/s41598-018-31951-z

19. Paltridge NG, Milham PJ, Ortiz-Monasterio JI Velu G, Yasmin Z, Palmer LJ et al. 2012; Energy-dispersive X-ray fluorescence spectrometry as a tool for zinc, iron and selenium analysis in whole grain wheat. Plant Soil 361:261–269 (2012). doi: 10.1007/s11104-012-1423-0

20. Alvarado G, Rodríguez FM, Pacheco A, Burgueño J, Crossa J, Vargas M et al. 2020; Meta-r: a software to analyze data from multi-environment plant breeding trials. The Crop Journal 8(5):745–756. doi: 10.1016/j.cj.2020.03.010

21. Poland JA, Brown PJ, Sorrells ME, Jannink JL. 2012; Development of high-density genetic maps for barley and wheat using a novel two-enzyme genotyping-by-sequencing approach. PLoS One 7(2):e32253. 10.1371/journal.pone.0032253

22. Glaubitz JC, Casstevens TM, Lu F, Harriman J, Elshire RJ, Sun Q, Buckler ES. 2014; TASSEL-GBS: a high capacity genotyping by sequencing analysis pipeline. PLoS One 9(2):e90346. doi: 10.1371/journal.pone.0090346

23. Kuzay S, Xu Y, Zhang J, Katz A, Pearce S, Su Z et al. 2019; Identification of a candidate gene for a QTL for spikelet number per spike on wheat chromosome arm 7AL by high-resolution genetic mapping. Theor Appl Genet 132(9):2689–2705. doi: 10.1007/s00122-019-03382-5

24. Shim H, Chasman DI, Smith JD, Mora S, Ridker PM, Nickerson DA et al. 2015; A multivariate genome-wide association analysis of 10 LDL subfractions, and their response to statin treatment, in 1868 Caucasians. PLoS One 10(4):e0120758. doi: 10.1371/journal.pone.0120758

25. Krishnappa G, Singh AM, Chaudhary S, Ahlawat AK, Singh SK, Shukla RB et al. 2017; Molecular mapping of the grain iron and zinc concentration, protein content and thousand kernel weight in wheat (Triticum aestivum L.). PloS one 12(4):e0174972. doi: 10.1371/journal.pone.0174972

26. Velu G, Ortiz-Monasterio I, Cakmak I, Hao Y, Singh RÁ. 2014; Biofortification strategies to increase grain zinc and iron concentrations in wheat. J Cereal Sci 59(3):365–372. doi: 10.1016/j.jcs.2013.09.001

27. Distelfeld A, Cakmak I, Peleg Z, Ozturk L, Yazici AM, Budak H, Saranga Y, Fahima T. 2007; Multiple QTL-effects of wheat Gpc-B1 locus on grain protein and micronutrient concentrations. Physiol Plant 129(3): 635–643. doi: 10.1111/j.1399-3054.2006.00841.x

28. Uauy C, Distelfeld A, Fahima T, Blechl A, Dubcovsky J. 2006; A NAC Gene regulating senescence improves grain protein, zinc, and iron content in wheat. Science 314(5803):1298–301. doi: 10.1126/science.1133649

29. Tong J, Zhao C, Sun M, Fu L, Song J, Liu D. 2022; High Resolution Genome Wide Association Studies Reveal Rich Genetic Architectures of Grain Zinc and Iron in Common Wheat (Triticum aestivum L.). Front Plant Sci 13:840614.doi: 10.3389/fpls.2022.840614

30. Bhatta M, Baenziger PS, Waters BM, Poudel R, Belamkar V, Poland J, Morgounov A. 2018; Genome-Wide Association Study Reveals Novel Genomic Regions Associated with Grain Minerals in Synthetic Hexaploid Wheat. Int J Mol Sci 19(10):3237. doi: 10.3390/ijms19103237

31. Sun M, Luo Q, Zheng Q, Tong J, Wang Y, Song J et al. 2023; Molecular characterization of stable QTL and putative candidate genes for grain zinc and iron concentrations in two related wheat populations. Theor Appl Genet 136(10):217. doi: 10.1007/s00122-023-04467-y

32. Chao S, Dubcovsky J, Dvorak J, Luo MC, Baenziger SP, Matnyazov R, Clark DR et al. 2010; Population-and genome-specific patterns of linkage disequilibrium and SNP variation in spring and winter wheat (Triticum aestivum L.). BMC Genomics 11:727. doi: 10.1186/1471-2164-11-727

33. Darrier B, Rimbert H, Balfourier F, Pingault L, Josselin AA et al. 2017; High-Resolution Mapping of Crossover Events in the Hexaploid Wheat Genome Suggests a Universal Recombination Mechanism. Genetics 206(3): 1373–1388. doi: 10.1534/genetics.116.196014

